# Synaptic mitochondrial oxidative stress drives individual variability in age-related cognitive decline in mice

**DOI:** 10.64898/2026.01.08.698304

**Authors:** Rui Yamada, Hirotaka Nagai, Chisato Numa, Yunhui Zhu, Midori Nagai, Kohei Ota, Yusuke Kawashima, Nobuhiko Ohno, Tomoyuki Furuyashiki

## Abstract

Aging leads to cognitive decline with considerable individual variability, yet the biological mechanisms remain unclear. Here we performed ultrastructural and proteomic analyses of the medial prefrontal cortex (mPFC) alongside behavioral assessments of attentional set shifting in mice across age groups. Although reduced synaptic density did not consistently lead to cognitive decline, proteomic analyses of synaptosomes and whole tissue revealed that the molecular signatures associated with individual variability in cognitive decline were distinct from those associated with chronological aging, and that synaptic mitochondria and their proteins were more abundant in aged mice with greater cognitive decline. Moreover, treatment with the mitochondria-targeted antioxidant MitoQ reduced the abundance of synaptic mitochondrial proteins, including pro-apoptotic proteins, and mitigated age-related cognitive decline. These findings demonstrate that synaptic mitochondrial oxidative stress in the mPFC, distinct from chronological age-related processes, contributes to individual variability in age-related cognitive decline and offers a potential target for prevention and intervention.

## Introduction

Aging is a complex biological process that impacts multiple brain systems and often leads to cognitive decline. Among the most affected regions is the prefrontal cortex (PFC), which plays a key role in higher-order cognitive functions such as working memory, attention, and behavioral flexibility. Age-related PFC dysfunction has been well documented in humans, non-human primates, and rodents, and is strongly associated with impairments in executive function and quality of life in late adulthood^1–3^. Cognitive flexibility is particularly sensitive to prefrontal aging and serves as a cross-species measure of executive control^4–9^.

Across species, aging is accompanied by dendritic atrophy and reduced synaptic density in PFC neurons, along with decreases in synaptic molecules^10,11^. Altered central metabolism and elevated oxidative stress are also reported in aged brains^12^. Mitochondrial dysfunction, a well-established hallmark of brain aging, is especially critical. Functional mitochondria at presynaptic terminals support synaptic transmission by supplying ATP and neurotransmitter precursors^13^. However, aging disrupts mitochondrial function, promoting oxidative stress through impaired oxidative phosphorylation and the release of proapoptotic molecules such as cytochrome c^14,15^. Thus, mitochondrial homeostasis must be maintained by removing damaged mitochondria through mitophagy to prevent these effects^16^. However, how these age-associated molecular and synaptic changes relate to cognitive decline remains elusive.

While cognitive decline is a common feature of aging, substantial individual variability exists^17^. Similar variability has been observed in rodent studies, providing valuable models for uncovering the molecular and synaptic mechanisms underlying this process^6,7^. In this study, we used the attentional set shifting test to assess individual variability in cognitive flexibility among aged mice and combined behavioral data with ultrastructural and proteomic analyses of the medial prefrontal cortex (mPFC), a key region for attentional set shifting. Our results showed that proteomic profiles linked to individual variability in cognitive aging are distinct from chronological age-related changes and feature a higher abundance of mitochondria and mitochondrial proteins at synapses for greater cognitive decline. Moreover, treatment with the mitochondria-targeted antioxidant MitoQ reduced the levels of synaptic mitochondrial proteins, including pro-apoptotic molecules, and restored cognitive flexibility in aged mice. Together, these findings demonstrate that synaptic mitochondrial dysfunction, distinct from chronological age-related processes, contributes to individual variability in age-related cognitive decline, offering a potential therapeutic target for cognitive aging.

## Methods

### Animals

Male and female C57BL/6J mice were obtained from Jackson Laboratory Japan (Yokohama, Japan), and male and female C57BL/6N mice were purchased from Japan SLC (Shizuoka, Japan). Mice were categorized by age as follows: young (8-12 weeks), middle-aged (50 weeks), and aged (75-85 weeks). All animals were housed in a controlled facility under stable temperature and humidity conditions with a 12-h light/12-h dark cycle. Food and water were provided ad libitum throughout the study, except when food deprivation was required for behavioral testing. Initially, mice were group-housed (4-5 per cage) and were transitioned to single housing prior to behavioral testing.

All experimental procedures were designed to minimize animal suffering and to reduce the number of animals used, in accordance with the principles of the 3Rs (Replacement, Reduction, and Refinement). Animal health and welfare were monitored daily, and humane endpoints were applied when necessary. All procedures were conducted in accordance with the ARRIVE guidelines and the NIH Guide for the Care and Use of Laboratory Animals and were approved by the Animal Care and Use Committees of Kobe University Graduate School of Medicine.

### Behavioral experiments

Prior to behavioral testing, mice were food-restricted to approximately 80% of their baseline body weight to enhance motivation for food rewards. All behavioral tests were conducted in operant conditioning chambers equipped with a touchscreen interface and an integrated food tray connected to an automatic pellet dispenser. Mice were first habituated to the operant chamber environment. During this phase, five food pellets (10 mg each) were placed directly on the food tray before the session, and mice were allowed to freely consume them. If any pellets remained uneaten, the same habituation procedure was repeated the following day. Once a mouse consumed all five pellets, it proceeded to the next training phase. To familiarize mice with the food dispenser mechanism, pellets were delivered automatically every 10 s over a 15-min period, totaling approximately 90 pellets. Mice were then allowed an additional 45 min to consume the pellets, resulting in a 60-min session. If pellets were left uneaten, the same procedure was repeated the next day to ensure proper acclimatization. In the next phase, visual stimuli consisting of horizontal or vertical lines were displayed on either side of the touchscreen. Touching either side of the screen with the nose resulted in the delivery of a food pellet. Each session lasted for one hour or until the mouse reached 150 screen touches, whichever occurred first. Mice that performed fewer than 30 nose pokes were re-exposed to the same test on the following day.

Following touchscreen habituation, mice underwent a visual discrimination test in which they were required to touch a specific stimulus (horizontal or vertical lines) to receive a food reward. In the experiments without fiber photometry (Figs. 1 and 4), sessions lasted one hour or until 100 screen touches were reached, whichever occurred first, and mice completed two sessions per day. In experiments combined with fiber photometry (Extended Data Fig. 2), sessions lasted one hour or until 150 screen touches were reached, whichever occurred first, and mice completed one session per day. Mice continued training daily until they achieved at least 80% correct responses in a single session or exceeded 75% correct responses in at least two sessions over eight training days. If performance criteria were not met, training was terminated after 42 sessions. To assess the attentional set shifting ability, mice then underwent a response direction test, where they were required to touch a specific side of the screen (left or right), irrespective of the visual stimulus, to receive a reward. Each session lasted one hour or until 100 screen touches were reached, and mice completed one session daily for five consecutive days. Two aged male C57BL/6N mice and one aged male C57BL/6J mouse were excluded from analysis, because they performed fewer than 10 trials in the visual discrimination test.

**Fig. 1.**
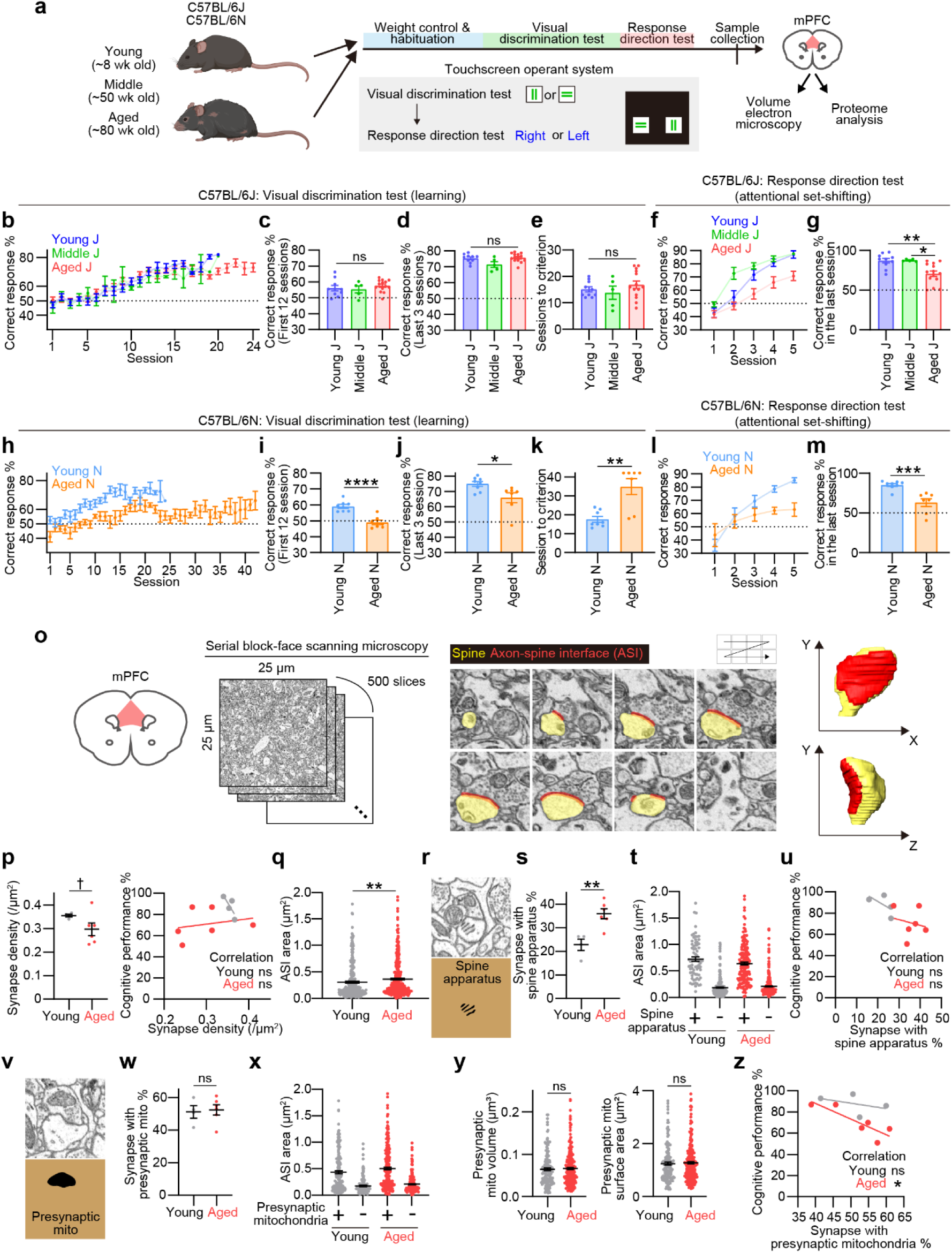
Aged C57BL/6J mice show individual variability in cognitive decline associated with presynaptic mitochondrial abundance in the mPFC. (a) Schematic overview of the touchscreen-based operant conditioning paradigm. (b-g) Behavioral performance of young, middle-aged, and aged C57BL/6J mice in the visual discrimination and response direction tests. For the visual discrimination test, the correct response rate averaged within each session throughout the experiment (b), over the first 12 sessions (c), and over the last 3 sessions (d), as well as the number of sessions required to reach the performance criterion (e), are shown. For the response direction test, the correct response rate within each trial throughout the experiment (f) and within the last session (g) are shown. Note that the correct response rate in the last session varies considerably among aged C57BL/6J mice. (h-m) Behavioral performance of young and aged C57BL/6N mice in the visual discrimination and response direction tests. For the visual discrimination test, the correct response rate averaged within each session throughout the experiment (h), over the first 12 sessions (i), and over the last 3 sessions (j), as well as the number of sessions required to reach the performance criterion (k), are shown. For the response direction test, the correct response rate within each session throughout the experiment (l) and within the last session (m) are shown. Note that the behavioral performance is uniformly impaired among aged mice in both tests. (o) Schematic illustration of volume electron microscopy used to examine synaptic ultrastructure and schematic of axon-spine interface (ASI) annotation, a structural proxy for synapse size. (p) (Left) Synapse density. Three regions within one image stack were analyzed per mouse; each data point represents one mouse. (Right) Correlation between synapse density and cognitive performance (measured by the last session of the response direction test). (q) ASI size. (r) Representative image of a spine apparatus, typically located at the neck of large spine heads. (s) Proportion of synapses with spine heads containing a spine apparatus. (t) ASI size for synapses with or without a spine apparatus. (u) Correlation between the proportion of spine apparatus-containing synapses and cognitive performance. (v) Representative image of presynaptic mitochondria. (w) Proportion of synapses containing presynaptic mitochondria. (x) ASI size for synapses with or without presynaptic mitochondria. (y) Volume and surface area of presynaptic mitochondria. (z) Correlation between the proportion of presynaptic mitochondria-containing synapses and cognitive performance. Scale bars: one edge of each image in (o), (r), and (v) corresponds to 2 μm. Data are presented as mean ± SEM. Statistical analyses were performed using one-way ANOVA with Holm-Sidak post hoc multiple comparisons (c,d,e,g), unpaired two-tailed Student’s *t*-test (i,j,k,m,p,q,s,w,y), and Pearson’s correlation test (p,u,z). † *P* < 0.1, * *P* < 0.05, ** *P* < 0.01, \*\*\**P* < 0.001, \*\*\*\**P* < 0.0001; ns, not significant. Ten young, five middle-aged, and 14 aged male C57BL/6J mice were used in (b-g). Eight young and seven aged male C57BL/6N mice were used in (h-m). Four young and six aged male C57BL/6J mice following behavioral tests were used in (o-z).

For pharmacological experiments, male C57BL/6J mice were provided with drinking water containing either MitoQ (mitoquinone mesylate, 100 μM) or decyl triphenylphosphonium bromide (dTPP) (100 μM), together with β-cyclodextrin (400 μM), from 55 weeks to 75 weeks and throughout the subsequent behavioral tests. The dosing regimen was based on previous studies^18,19^. dTPP is structurally identical to MitoQ but lacks the ubiquinone moiety and thus serves as a control compound that retains the lipophilic cationic carrier without antioxidant functionality. Two MitoQ-treated mice were excluded from analysis, because they did not reach the performance criteria described above in the visual discrimination test.

A subset of the behavioral data from aged and young male C57BL/6J mice are reported elsewhere^20^.

### Fiber photometry

To monitor neuronal activity in the medial prefrontal cortex (mPFC), fiber photometry was employed. Mice were first deeply anesthetized with isoflurane and placed in a stereotaxic apparatus. A total volume of 500 nL of artificial cerebrospinal fluid (124 mM NaCl, 3 mM KCl, 26 mM NaHCO₃, 2 mM CaCl₂, 1 mM MgSO₄, 1.25 mM KH₂PO₄, and 10 mM D-glucose) containing adeno-associated viral (AAV) vectors encoding the calcium indicator GCaMP7c under the human synapsin 1 promoter (made from pGP-AAV-syn-jGCaMP7c variant 1513-WPRE; titer: 1.0-2.5 × 10¹³ GC/mL) was injected into the left mPFC (coordinates: AP +1.8 mm, ML +0.4 mm, DV –2.8 mm from bregma). pGP-AAV-syn-jGCaMP7c variant 1513-WPRE was a gift from Douglas Kim & GENIE Project (Addgene plasmid #105321; http://n2t.net/addgene:105321; RRID:Addgene_105321)^21^. AAV vectors were created as previously described^22,23^. Following injection, an optical fiber (200 μm core diameter, 0.5 NA, multimode, high OH, Thorlabs Inc.), secured within a 6.4 mm ceramic ferrule, was implanted 0.2 mm above the injection site and fixed to the skull using dental cement (G-SEM ONE Neo; GC Corporation, Tokyo, Japan). Mice were housed individually and allowed to recover for three weeks before behavioral testing.

Fiber photometry recordings were performed based on established protocols with minor modifications^24,25^. Two excitation wavelengths were delivered via an optical fiber: 465 nm (Ca²⁺-dependent excitation of GCaMP7c) and 405 nm (isosbestic control for motion artifacts). The 465 nm LED was modulated at 572.205 Hz and the 405 nm LED at 208.616 Hz. Light intensities were adjusted to maintain raw input signals between 1-4 V and lock-in demodulated signals between 0.1-0.9 V at baseline. Fluorescence emission was collected by a photoreceiver (Newport Corporation, Irvine, CA, USA) and transmitted to the fiber photometry console for digitization. Δ F/F was calculated from calcium-dependent and independent signals using Doric Neuroscience Studio.

During recordings, 5 V TTL pulses were sent from the operant task chamber to the photometry system to enable time-locked analyses. Two behavioral events were analyzed: (1) correct response (touching the correct side of the touchscreen), and (2) error response (touching the incorrect side). For each event, fluorescence signals from a 10-s window centered on the event were extracted. The ΔF/F traces were smoothed using a 0.5-s moving average. Outlier trials were excluded if the mean ΔF/F within the 10-s window for a given trial deviated by more than ±2 SD from the overall mean across all trials.

### Sample collection

After the behavioral experiments, mice were placed in their home cages with food and water available ad libitum for 2-3 weeks. mPFC tissues were collected for either proteomic analysis or volume electron microscopy.

For proteomic analysis, mice were deeply anesthetized via intraperitoneal injection of sodium pentobarbital (100 mg/kg; Nacalai Tesque, Kyoto, Japan), followed by transcardial perfusion with ice-cold saline to remove blood. The mPFC, including the anterior cingulate cortex, prelimbic cortex, and infralimbic cortex, was rapidly dissected in the synaptosome isolation buffer, containing 320 mM sucrose, 5 mM HEPES, 0.05 mM dithiothreitol (DTT), 0.1 mM phenylmethylsulfonyl fluoride, 2 U/mL RNaseOUT, and a complete protease inhibitor cocktail (pH 7.4). Tissue was homogenized using a Dounce homogenizer. A portion of the homogenate was reserved for whole fraction analysis. The remaining homogenate was centrifuged at 1,000 × g for 10 min at 4°C to remove nuclei. The supernatant was then centrifuged at 20,400 × g for 10 min at 4°C to eliminate microsomal components. The resulting pellet was resuspended in the synaptosome isolation buffer and layered onto a discontinuous Percoll gradient (15% and 23%). Ultracentrifugation was performed at 31,000 × g for 8 min at 4°C. The synaptosome-enriched fraction was carefully collected and processed for proteomic analysis.

For volume electron microscopy, mice were anesthetized with sodium pentobarbital and perfused transcardially with pre-warmed saline (37°C), followed by half Karnovsky fixative containing 2.5% glutaraldehyde, 2% paraformaldehyde, and 2 mM calcium chloride in 0.1 M cacodylate buffer (pH 7.4). Brains were removed and post-fixed overnight in the same fixative at 4°C. The following day, 120 μm-thick coronal sections were prepared using a vibratome in the cutting solution (0.1 M cacodylate buffer, pH 7.4, with 2 mM calcium chloride). Brain slices were stored at −30°C in cryoprotectant until further processing.

### Volume electron microscopy

The samples were processed as previously described with some modifications^26,27^. Briefly, the infralimbic cortex was punched out using a 1.5-mm trepan biopsy tool in the cryoprotectant solution. The punched-out specimen was washed in the cutting solution for 3 min four times at 4℃, incubated for 1 h on ice with a solution of reduced osmium solution containing 1.5% potassium ferrocyanide, 2% osmium tetroxide, and 2mM calcium chloride in 0.1M cacodylate buffer, and exposed to a solution of 1% thiocarbohydrazide for 20 min at room temperature. The specimen was then incubated in 2% osmium tetroxide for 30 min at room temperature and incubated overnight with 1% uranyl acetate at 4℃ overnight. Then the specimen was stained with a solution of lead aspartate for 30 min at 60℃ (pH 5.5) and dehydrated using ice-cold solutions of freshly prepared 20%, 50%, 70%, 90%, 100%, and 100% anhydrous ethanol for 5 min each, followed by ice-cold anhydrous acetone for 10 min twice. The specimens were then placed in the solution containing 50% Quetol 812 resin and 50% acetone for 45 min, and then 100% Quetol 812 resin for 45 min 3 times at 35℃. To increase electroconductivity of resin, we used carbon black filler, Ketjen black^28^. Ketjen black was mixed with Quetol 812 resin as 7:93 ratio. The specimen was flat embedded with ACLAR embedding film and kept in a 70℃ oven for 72 h.

The resin-embedded specimen was observed by Sigma scanning electron microscope (Carl Zeiss Microscopy, Goettingen, Germany) equipped with 3View technology (Gatan, Inc, Pleasanton, CA, USA). In serial block-face scanning electron microscopy (SBEM), serial images were obtained by scanning the face of an unsliced block of tissue placed inside the microscope, then cutting off ultrathin slices using an automated microtome within the instrument. The newly exposed surface of the sliced block was rescanned until a stack of images was obtained. Serial images were acquired using an aperture of 30μm, high vacuum, acceleration voltage of 1.5kV, image size of 5,000 by 5,000 pixels, and image resolution (xy plane) of 5 nm, while ultrathin sections were cut at a nominal thickness of 50 nm. A stack of ∼500 images was acquired per mouse (15,625 μm³), in layer 2/3 of the infralimbic cortex. Images were Gaussian filtered and automatically aligned using the open-source software Fiji.

For ultrastructural image analysis, regions densely filled with neuropils were cropped to a volume of 10 μm × 10 μm × 10 μm. From each image stack, 80 synapses were randomly selected and analyzed using the TrakEM2 plugin within Fiji software. The axon-spine interface (ASI) area was quantified as a structural index of synapse size. To define this interface, dendritic spines were annotated, and the surface area in direct contact with the apposed axon forming a synapse was identified. The interface area was then calculated using Amira software (Thermo Fisher Scientific, Waltham, MA, USA). In addition to synapse size, several ultrastructural features were assessed as histological parameters. Specifically, the presence or absence of presynaptic mitochondria and the presence or absence of a spine apparatus within the dendritic spine were recorded. For a subset of ∼30 randomly selected synapses per stack, the presence of tripartite synapses, where astrocytic processes contact the synaptic cleft, was evaluated. The proportion of the ASI perimeter covered by astrocytic processes was calculated, based on previously established methods^29^.

### Proteomic analysis

Proteomic sample preparation and analysis were performed according to established protocols with minor modifications^30^. Samples were cleaned with cold acetone and incubated at −20°C for 2 h. After centrifugation, precipitates were dissolved in 100 mM Tris (pH 8.5) with 0.5% sodium dodecanoate using an ultrasonic homogenizer. Protein concentration was measured using a BCA assay and adjusted to 1 µg/µL. Disulfide bonds were reduced with 10 mM DTT at 50°C for 30 min, followed by cysteine alkylation with 30 mM iodoacetamide (IAA) in the dark at room temperature for 30 min. The reaction was quenched with 60 mM cysteine. For protein digestion, 150 µL of 50 mM ammonium bicarbonate was added, and proteins were digested overnight at 37°C with Lys-C and trypsin (400 ng each). Peptides were acidified with 5% trifluoroacetic acid, centrifuged, desalted using a C18 spin column, dried, and reconstituted in 3% acetonitrile (ACN) with 0.1% formic acid. Peptide concentration was adjusted to 500 ng/µL.

For nanoLC-MS/MS analysis, 500 ng of peptides were injected into an UltiMate 3000 RSLCnano LC System (Thermo Fisher Scientific) with an Aurora Series emitter column (75 µm × 250 mm) at 60°C. Solvents were 0.1% formic acid in water (A) and 0.1% formic acid in 80% ACN (B). The gradient was 2-4% B for 0-3 min, 4-34% B for 3-113 min, 34-65% B for 113-118 min, and 65% B for 118-123 min at 400 nL/min for 0-3 min and 200 nL/min for 3-123 min. The Orbitrap Exploris 480 MS (Thermo Fisher Scientific) was used in ESI positive mode. Full MS1 scans (495-745 m/z) alternated with DIA MS2 scans (200-1800 m/z) using a resolution of 15,000 for MS1 and 45,000 for MS2, AGC target of 3e6, and isolation window of 4.0 m/z.

Data were analyzed using Scaffold DIA (Proteome Software) with the Mouse UniProtKB/Swiss-Prot database. Parameters included HCD fragmentation, 10 ppm precursor and fragment tolerance, staggered DIA acquisition, trypsin digestion, 2-4 peptide charge, max one missed cleavage, and carbamidomethylation [C] as a fixed modification. Both peptide and protein FDR were set below 1%. Quantification results were then normalized to the median value of each sample. Proteins that were detected in only one sample were excluded from analysis.

The expression levels of proteins were represented as the signal intensities obtained from the nanoLC-MS/MS system or as their Z-score normalized value to enable the comparison of multiple proteins in the same analyses. To identify differentially-expressed proteins depending on age (Age DEPs), a one-way ANOVA was performed to compare the abundance of each protein across young, middle-aged, and aged groups. Proteins with *P* < 0.05 were defined as Age DEPs. To identify differentially-expressed proteins depending on cognitive performance (Score DEPs), Pearson correlation analyses were performed to examine the relationship between protein abundance and behavioral accuracy among mice in the aged group. Proteins with *P* < 0.05 were defined as Score DEPs. Hierarchical clustering analyses were then performed using Age DEPs and Score DEPs, and the major clusters were defined as C1 and C2 in Fig.2.

**Fig 2.**
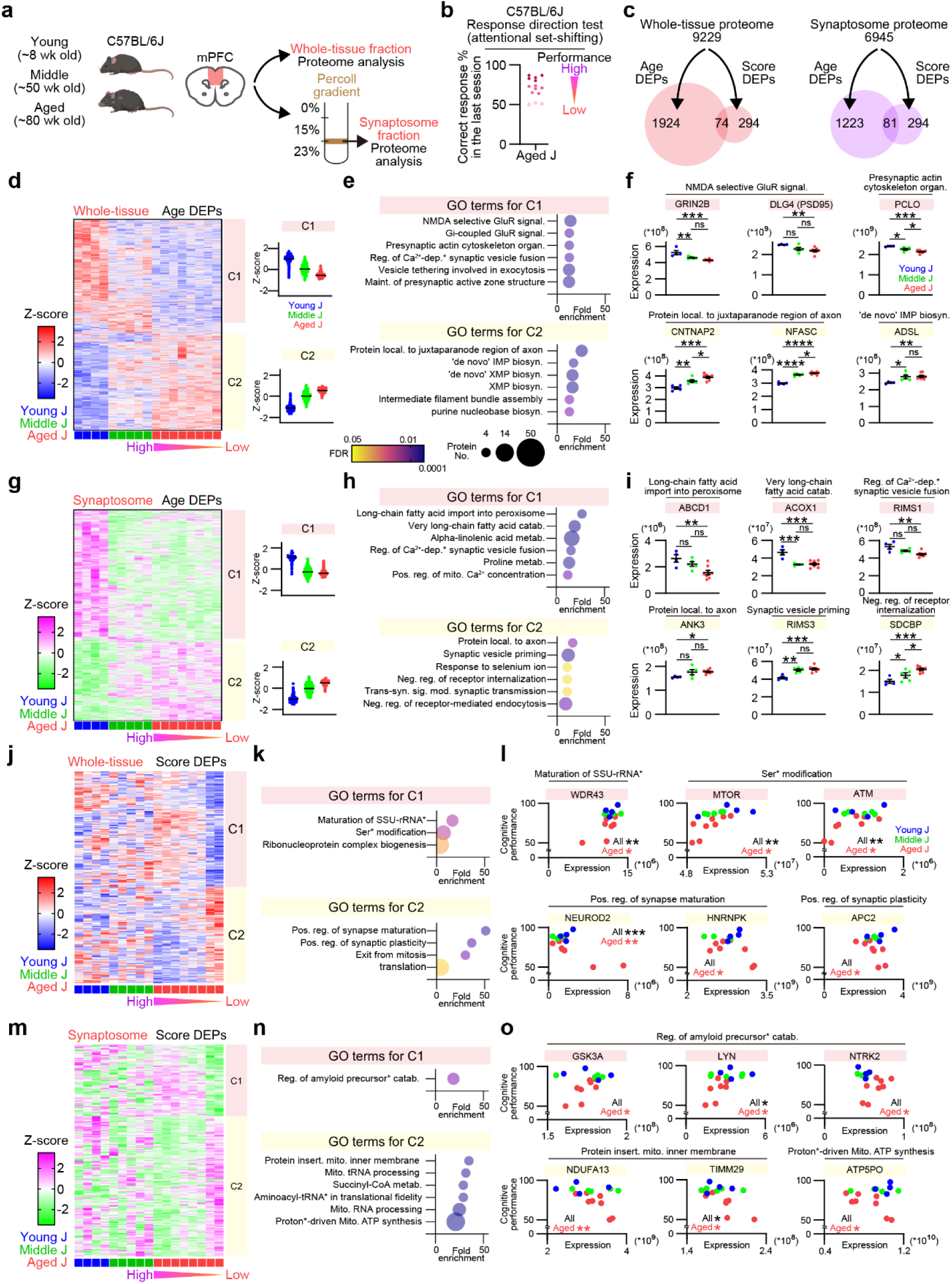
The mPFC proteome reveals distinct molecular signatures associated with chronological aging and individual variability in cognitive decline. (a) Schematic of sample preparation for proteomic analyses. Whole tissue and synaptosome fractions were isolated from the mPFC of male C57BL/6J mice across different age groups using a density gradient centrifugation method. (b) Cognitive performance during the last session of the response direction test in aged C57BL/6J mice. Data from aged mice in Fig. 1g are shown again to illustrate individual variability in cognitive decline. (c) Age DEPs and Score DEPs identified in whole tissue and synaptosome fractions (see Methods), with the overlap between Age DEPs and Score DEPs at the chance level. (d-i) Clustering analysis of Age DEPs in whole tissue (d-f) or synaptosome (g-i) samples, identifying two clusters, Cluster 1 (downregulated in aged mice) and Cluster 2 (upregulated in aged mice) (d,g), their enriched GO terms (e,h), and the expression of representative proteins (f,i). (j-o) Clustering analysis of Score DEPs in whole tissue (j-l) or synaptosome (m-o) samples, identifying two clusters, Cluster 1 (downregulated in aged mice with greater cognitive decline) and Cluster 2 (upregulated in aged mice with greater cognitive decline) (j,m), their enriched GO terms (k,n), and the expression of representative proteins (l,o). Abbreviations: signal.; signaling pathway, dep.*; dependent activation of, maint.; maintenance, local.; localization, biosyn.; biosynthetic process, reg.; regulation, pos.; positive, metab.; metabolic process, mito.; mitochondrial, neg.; negative. Data are presented as mean ± SEM. Statistical analyses were performed using one-way ANOVA with Holm-Sidak post hoc multiple comparisons (f,i) and Pearson’s correlation test (l,o). \**P* < 0.05, \*\**P* < 0.01, \*\*\**P* < 0.001, \*\*\*\**P* < 0.0001; ns, not significant. Four young, five middle-aged, and eight aged male C57BL/6J mice were used following behavioral tests.

Gene ontology (GO) analysis was performed on designated proteins using the publicly available resource at https://geneontology.org/. GO terms containing more than three and fewer than 500 assigned proteins were included in the analysis. Up to six out of the top ten GO terms based on fold enrichment were displayed. When calculating log₂ fold changes between groups, if a protein was undetected in one group, a placeholder value of +10 or −10 was assigned for visualization purposes.

Protein-protein interaction networks were constructed using Cytoscape software^31^, referencing interaction data from the STRING database. Proteins with a high number of connections (i.e., high degree centrality) within the network were defined as hub proteins. Proteins constituting respiratory chain complexes were identified using publicly available database (https://www.genenames.org/).

### Statistical analyses

Data are shown as means ± SEM. Statistical analyses were performed using Prism 10.6 software (GraphPad Software, San Diego, CA, USA). *P* values less than 0.05 were considered statistically significant. Comparisons between two groups were analyzed using an unpaired *t*-test (Fig. 1i-k,m,p,q,s,w,y,4b,c,h,j,k). Comparisons among more than two groups were analyzed using one-way ANOVA followed by Holm-Sidak’s post hoc multiple comparison tests (Fig. 1c-e,g,2f,i). Correlation analyses were performed by Pearson’s correlation test (Fig. 1u,z,2l,o,3e,f,4f). Comparisons against a designated value were analyzed using a one-sample *t*-test (Fig. 3g,4g; compared with 0). See the figure legends for details on the statistical analyses used in the Extended Data Figs.

**Fig 3.**
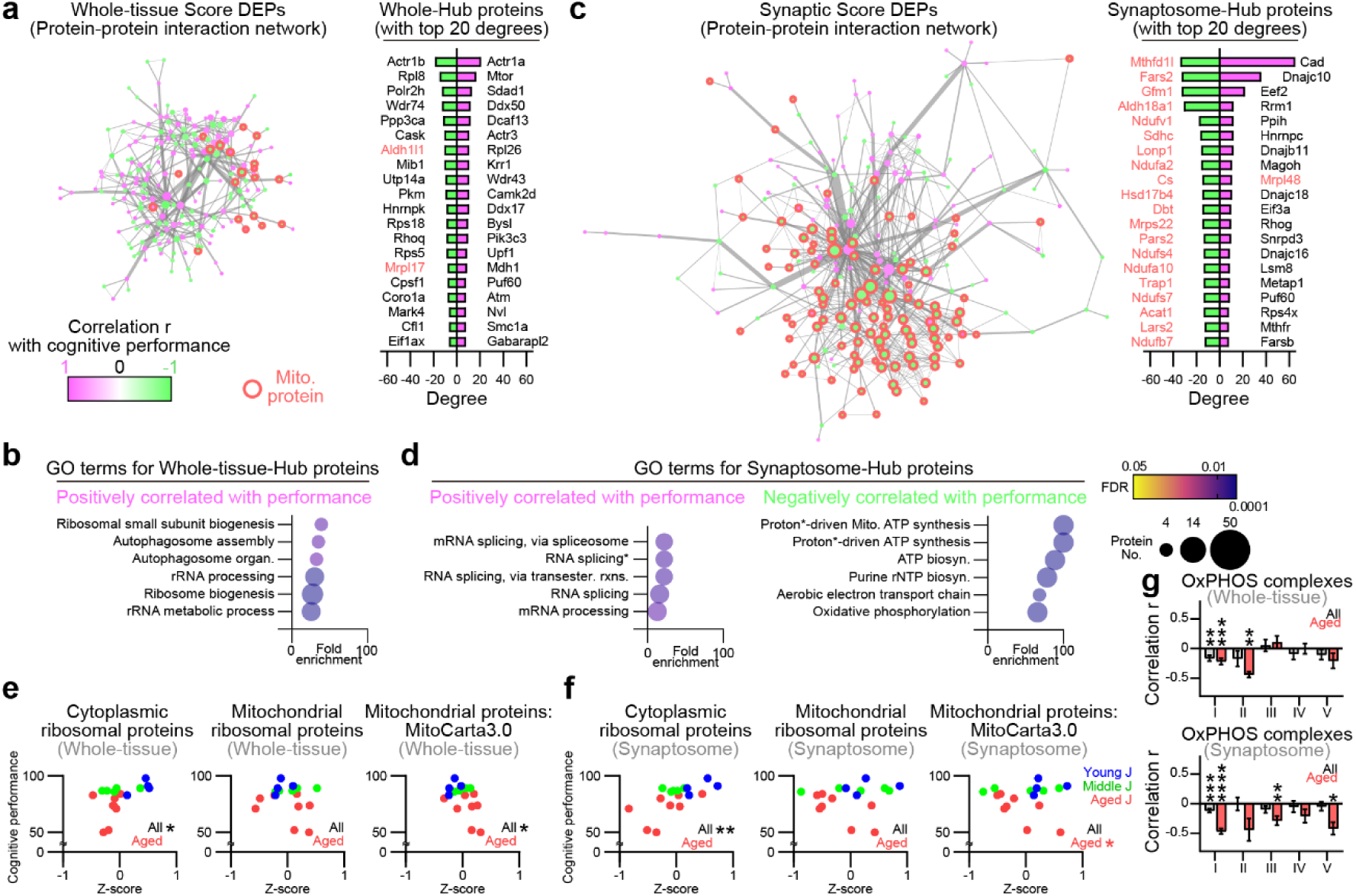
Synaptic mitochondrial proteins are more abundant in aged mice with greater cognitive decline. (a-d) Protein-protein interaction networks and the top 20 hub proteins ranked by degrees (number of connections for each node) for Score DEPs in whole tissue (a) and synaptosome (c) fractions, along with the GO terms associated with these hub proteins (b and d, respectively). Abbreviations: organ.; organization, splicing*; splicing, via transesterification reactions with bulged adenosine as nucleophile, transester.; transesterification, rxns.; reactions, proton*; proton motive force, biosyn.; biosynthetic process. (e,f) Correlations between the expression levels of cytoplasmic or mitochondrial ribosomal proteins, or total mitochondrial proteins in whole tissue (e) and synaptosome (f) fractions and cognitive performance (measured by the last session of the response direction test). (g) Average correlation coefficients between the expression levels of proteins in each respiratory chain complex and cognitive performance. Statistical analyses were performed using Pearson’s correlation tests (e-g) followed by one-sample *t*-test against the chance level (0) in (g). Data are presented as mean ± SEM. \*\**P* < 0.01, \*\*\**P* < 0.001, \*\*\*\**P* < 0.0001. Four young, five middle-aged, and eight aged male C57BL/6J mice were used following behavioral tests.

## Results

### Aged C57BL/6J mice show individual variability in cognitive decline

To evaluate cognitive aging, we assessed cognitive flexibility using an attentional set-shifting test conducted in touchscreen-based operant chambers (Fig. 1a). Mice were first trained on a visual discrimination test, in which two distinct visual stimuli (vertical or horizontal lines) were simultaneously presented on a touchscreen. They had to select the predetermined correct stimulus, regardless of its position (left or right), to receive a food reward. After reaching a criterion accuracy, mice advanced to a response-direction test, where they were required to choose the correct side (left or right), irrespective of the stimulus pattern, to obtain the reward.

While C57BL6 mice are widely used in behavioral research, two substrains, C57BL/6J and C57BL/6N, reportedly show distinct age-related behavioral abnormalities in exploration, anxiety, and sociability^32^. We therefore compared male mice of these substrains across three age groups: young (8-12 weeks), middle-aged (50 weeks), and aged (75-85 weeks). All C57BL/6J mice successfully learned the visual discrimination phase (Fig. 1b-e). In the response-direction phase, young and middle-aged mice adapted successfully; however, some aged mice failed, while others performed comparably to young mice, indicating individual variability in attentional set shifting ability (Fig. 1f,g). In contrast, whereas young C57BL/6N mice correctly performed the visual discrimination and response direction tests, aged C57BL/6N mice showed poor performance in both tests even after extended training, suggesting broader learning deficits (Fig. 1h-m). Male and female C57BL/6J mice showed similar age-related cognitive decline (Extended Data Fig. 1a-f), leading us to focus on males in subsequent experiments.

To examine neural correlates, we conducted fiber photometry recordings from the mPFC during task execution. In visual discrimination, aged but not young mice showed learning-dependent increases in mPFC activity after behavioral responses (Extended Data Fig. 2a,b). Similarly, during the response direction test, aged but not young mice showed strong mPFC activation in the early learning phase (Extended Data Fig. 2c,d). These activations occurred only during correct trials, indicating behavioral relevance and suggesting that aged mice rely on mPFC engagement for both visual discrimination learning and attentional set shifting.

### Age-related cognitive decline correlates to presynaptic mitochondrial abundance

To identify structural correlates of cognitive flexibility, as measured by final behavioral accuracy in the response direction test for attentional set shifting, we performed volume electron microscopy of the mPFC in aged and young mice (Fig. 1o). Aged mice showed a reduced density of synapses on dendritic spines, accompanied by a larger average axon-spine interface (ASI) and a higher proportion of spine apparatus-containing synapses, which were associated with large ASI areas, suggesting a preferential loss of smaller synapses lacking spine apparatuses (Fig. 1p-t). Despite these age-associated changes in synaptic density and structure, none of these parameters correlated with individual variability in cognitive performance (Fig. 1p,u). Astrocytic coverage of synapses, a component of the tripartite synapse, did not differ significantly between young and aged mice, as measured by the proportion of synaptic perimeters covered by astrocytic processes (19.34 ± 2.27% and 16.93 ± 1.45%, respectively; *P* = 0.35, Student’s *t*-test), nor did it correlate with cognitive flexibility in aged mice (Pearson’s correlation coefficient, r = 0.146; *P* = 0.7826). Notably, presynaptic mitochondria, typically observed in presynaptic terminals apposing dendritic spines with larger ASIs, showed a distinct relationship to age-related cognitive decline. Although neither the overall frequency nor morphology of presynaptic mitochondria, nor their associtation with ASI size, changed significantly with age (Fig. 1v-y), the proportion of presynaptic mitochondria was inversely correlated with cognitive flexibility in aged but not young mice (Fig. 1z). These findings suggest that reduced synaptic density in the mPFC of aged mice does not necessarily lead to cognitive decline, and that presynaptic mitochondrial abundance may underlie this individual variability.

### mPFC proteome reveals distinct signatures of aging and cognitive decline

To identify molecular correlates of age-dependent cognitive decline, we performed proteomic profiling of whole-tissue and synaptosome fractions in the mPFC, identifying proteins whose abundance varied across age groups (young, middle-aged, and aged) and correlated with attentional set shifting performance in aged mice (hereafter Age DEPs and Score DEPs, respectively) (Fig. 2a-c). Notably, the overlap between Age DEPs and Score DEPs was at the chance level (expected vs. observed overlap = 76.7 vs. 74 for whole tissue, *P* = 0.61; 70.4 vs. 81 for synaptosomes, *P* = 0.23; Fisher’s exact test), suggesting that neural mechanisms determining individual variability in cognitive decline are distinct from chronological aging processes.

To gain functional insights into Age DEPs and Score DEPs, we performed gene ontology (GO) analysis. In whole tissue Age DEPs, those decreasing with age were associated with NMDA receptor-mediated signaling, including GRIN2B and DLG4 (PSD95), and presynaptic actin cytoskeleton organization, including PCLO, whereas those increasing with age were associated with protein localization to the juxtaparanodal region and de novo inosine monophosphate (IMP) biosynthesis (Fig. 2d-f). In synaptosomal Age DEPs, those decreasing with age were associated with peroxisomal fatty acid catabolism, including ABCD1-3 and ACOX1, whereas those increasing with age were associated with axonal protein localization and synaptic vesicle priming (Fig. 2g-i). Thus, distinct age-related proteomic changes occurred in whole tissue and synaptosome fractions of the mPFC. In whole-tissue Score DEPs, proteins positively correlated with cognitive performance were associated with small subunit (SSU) rRNA maturation and serine/threonine kinase signaling, including MTOR and ATM, whereas those negatively correlated with cognitive performance were associated with positive regulation of synapse maturation and plasticity (Fig. 2j-l). In synaptosomal Score DEPs, proteins positively correlated with cognitive performance were associated with regulation of amyloid precursor protein catabolism, including GSK3A, LYN, NTRK2 (TrkB), whereas those negatively correlated were associated with mitochondrial processes, including mitochondrial protein regulation and metabolism (Fig. 2m-o).

### Synaptic mitochondrial proteins are more abundant in aged mice with greater cognitive decline

To identify the core protein signatures underlying individual variability in age-related cognitive decline, we examined the protein-protein interaction network of Score DEPs using the STRING database and identified hub proteins based on degrees (number of connections for each node). In the whole-tissue proteome, hub proteins positively correlated with cognitive performance were associated with ribosomal and autophagosomal regulation (Fig. 3a,b). In contrast, in the synaptosomal proteome, hub proteins positively correlated with cognitive performance were associated with mRNA splicing pathways, including HNRNPC, a splicing factor that also regulates mRNA metabolism and ribonucleoprotein assembly in the cytoplasm^33^, whereas those negatively correlated with cognitive performance were associated with mitochondrial oxidative phosphorylation (Fig. 3c,d).

Because the interaction network of synaptosomal Score DEPs was dominated by mitochondrial proteins, we focused subsequent analyses on mitochondrial proteins. In aged mice, mitochondrial proteins listed in the MitoCarta3.0 database, especially in synaptosomes, showed negative correlations with cognitive performance (Fig. 3e,f), which were absent in young and middle-aged mice. These negatively correlated proteins were involved in oxidative phosphorylation, the TCA cycle, and mitochondrial ribosomes (Fig. 3f,g). As synaptic mitochondria mainly originate from presynaptic terminals^34^, these findings indicate the association between presynaptic mitochondrial protein abundance and age-related cognitive decline. Conversely, cytoplasmic ribosomal proteins in synaptosomes were positively correlated with cognitive performance, suggesting differential regulation of cytoplasmic and mitochondrial translation. Such changes observed in synaptosomes were marginal in the whole tissue.

Collectively, these findings demonstrate that the synaptic mitochondrial proteins in the mPFC, likely derived from presynaptic terminals, are more abundant in aged mice with greater cognitive decline.

### MitoQ reduces synaptic mitochondrial protein abundance, preventing age-related cognitive decline

Given the negative correlations of the synaptic mitochondrial proteome with cognitive performance in aged mice, we hypothesized that synaptic mitochondrial dysfunction might underlie age-related cognitive decline. To test this hypothesis, we assessed the behavioral effects of MitoQ, a mitochondria-targeted antioxidant with beneficial effects in animal models of various diseases involving mitochondrial dysfunction^18,19,35^. Systemic MitoQ, but not its inactive analog dTPP, administered for 20 weeks significantly improved attentional set shifting (i.e., the response direction test) in aged mice without altering preceding visual discrimination learning (Fig. 4a-c), implicating mitochondrial dysfunction in age-related cognitive decline.

**Fig. 4.**
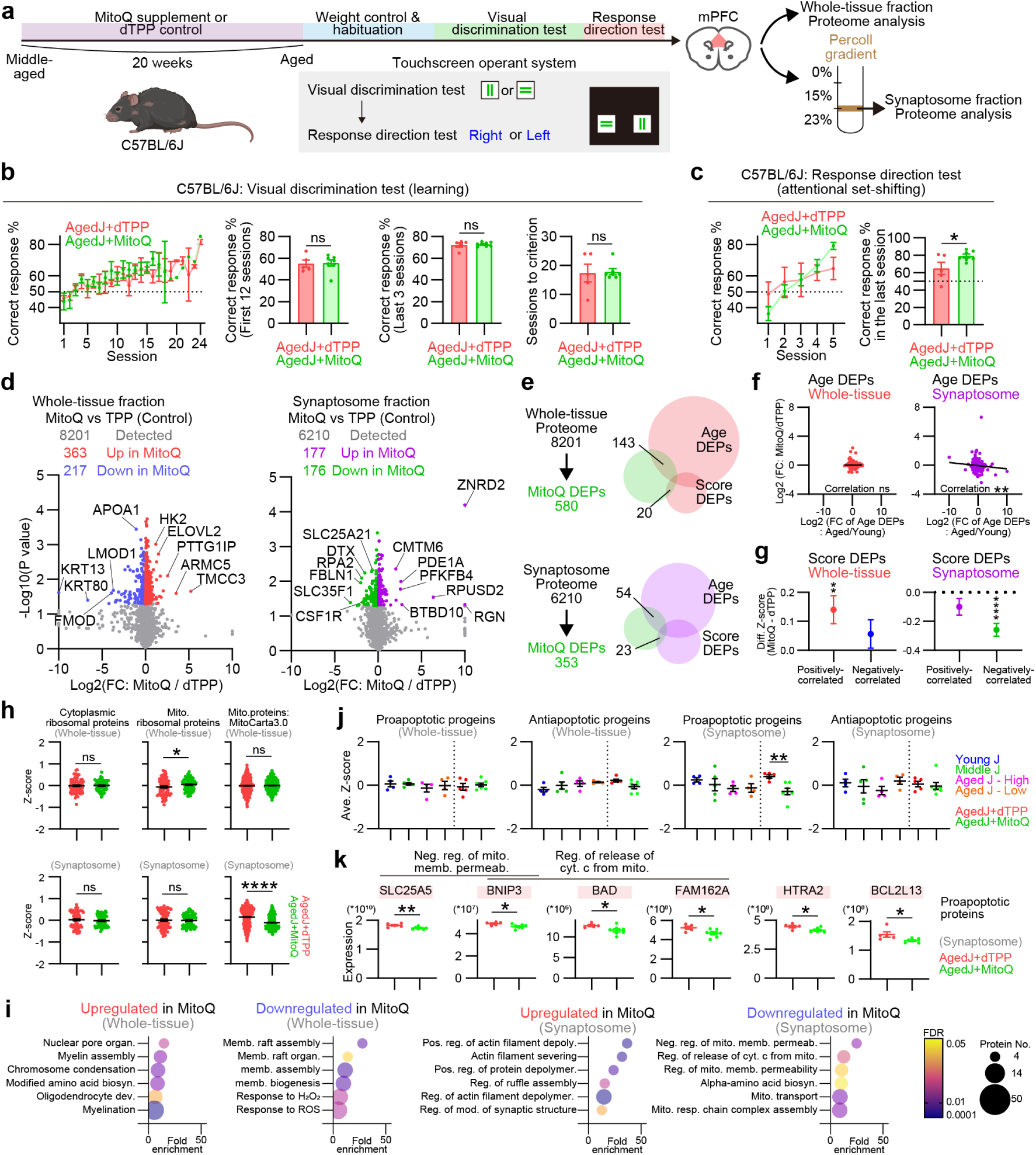
Treatment with the mitochondria-targeted antioxidant MitoQ reduces synaptic mitochondrial protein abundance, including pro-apoptotic proteins, and mitigates age-related cognitive decline. (a) Experimental timeline. C57BL/6J mice were administered either 100 µM MitoQ or dTPP (control) via drinking water from 55 to 75 weeks of age (20 weeks) and throughout the subsequent behavioral experiments, followed by proteomic analyses of whole tissue and synaptosome fractions of the mPFC. (b,c) Behavioral performance of MitoQ- or dTPP-treated mice in the visual discrimination and response direction tests. For the visual discrimination test, the correct response rate averaged within each session throughout the experiment, over the first 12 sessions, and over the last 3 sessions, as well as the number of sessions required to reach the performance criterion, are shown (b). For the response direction test, the correct response rate within each session throughout the experiment and within the last session are shown (c). (d) Volcano plots of MitoQ DEPs (see Results) in the whole tissue (left) and synaptosome (right) fractions. For visualization purposes, log_2_(fold change (FC)) values more than 10 or less than −10 are shown as 10 or −10, respectively. (e) Venn diagrams of MitoQ DEPs, Age DEPs, and Score DEPs in whole tissue and synaptosome fractions. (f) Correlations between age-associated changes (aged versus young mice) and MitoQ-induced changes (MitoQ- versus dTPP-treated mice) of Age DEPs in whole tissue and synaptosome fractions. (g) MitoQ-induced changes in the expression levels of Score DEPs in whole tissue and synaptosome fractions that are positively or negatively correlated with cognitive performance. (h) Expression levels of cytoplasmic (left) and mitochondrial (middle) ribosomal proteins, and total mitochondrial proteins defined by the MitoCarta3.0 database (right) in whole tissue (upper) and synaptosome (lower) fractions. (i) GO analysis of MitoQ DEPs of whole tissue and synaptosome fractions. Abbreviations: biosyn.; biosynthetic process, dev.; development, memb.; membrane, organ.; organization, pos.; positive, reg.; regulation, deploy.; depolymerization, mod.; modification, neg.; negative, cyt.; cytochrome, mito.; mitochondria, resp.; respiratory. (j) Averaged expression levels of pro- and anti-apoptotic proteins across age groups and in MitoQ-treated mice. (k) Expression of representative pro-apoptotic proteins in synaptosomes from MitoQ- and dTPP-treated aged mice. Data are presented as mean ± SEM. Statistical analyses were performed using unpaired two-tailed Student’s *t*-test (b,c,h,j,k), Pearson’s correlation test (f) and one-sample *t*-test (g). \**P* < 0.05, \*\*\**P* < 0.001, \*\*\*\**P* < 0.0001; ns, not significant. Five and seven mice that received drinking water containing dTPP and MitoQ, respectively, were used.

To investigate the mechanisms, we profiled MitoQ-regulated proteins (MitoQ DEPs) in mPFC whole tissue and synaptosomes (Fig. 4d). MitoQ DEPs partially overlapped with both Age DEPs and Score DEPs (Fig. 4e). In synaptosomes, but not in whole tissue, age-associated changes were negatively correlated with MitoQ-induced changes, indicating that MitoQ attenuated age-related alterations in the synaptosomal proteome (Fig. 4f). MitoQ also upregulated Score DEPs positively correlated with cognitive performance in whole tissue and downregulated those negatively correlated in synaptosomes (Fig. 4g), supporting a role for MitoQ-sensitive mitochondrial dysfunction in both chronological aging and individual variability in cognitive decline.

We next examined MitoQ-regulated processes related to cognitive decline. MitoQ downregulated mitochondrial proteins listed in the MitoCarta3.0 database selectively in synaptosomes and not in whole tissue (Fig. 4h), which were negatively correlated with cognitive performance as described above. MitoQ did not alter mitochondrial ribosomal proteins in synaptosomes, implicating other mitochondrial pathways. GO analysis of MitoQ DEPs showed that downregulated synaptosomal proteins were enriched for mitochondrial apoptotic processes, including negative regulation of mitochondrial membrane permeabilization and cytochrome c release (Fig. 4i). Consistent with this, MitoQ selectively decreased pro-apoptotic, but not anti-apoptotic, proteins in synaptosomes (Fig. 4j,k). GO terms also indicated effects beyond mitochondria: upregulation in synaptosomes of actin-reorganization factors (e.g., cofilin and ADF); upregulation in whole tissue of nuclear organization and myelination-related proteins; and downregulation in whole tissue of proteins linked to membrane raft assembly and responses to ROS, consistent with MitoQ’s antioxidant action.

Together, these data demonstrate that synaptic mitochondrial dysfunction underlies individual variability in age-related cognitive decline and can be ameliorated by a mitochondria-targeted antioxidant, possibly through multiple downstream pathways.

## Discussion

Aging is associated with cognitive decline that considerably varies among individuals, yet the biological mechanisms remain elusive. In this study, we performed ultrastructural and proteomic analyses of the mPFC together with behavioral assessments of attentional set shifting in mice across age groups. While not all aged mice with reduced synaptic density exhibit cognitive decline, synaptic mitochondria and their proteins, most likely derived from presynaptic terminals, are more abundant in aged mice with greater cognitive decline. Proteomic profiling of synaptosomes and whole tissue further shows that the neural mechanisms underlying individual variability in cognitive decline differ from those associated with chronological aging, each reflected by distinct protein-expression patterns. The role of synaptic mitochondrial dysfunction in age-related decline is further supported by the observation that treatment with the mitochondria-targeted antioxidant MitoQ reduced the synaptic mitochondrial protein abundance and mitigated cognitive decline. Thus, synaptic mitochondrial dysfunction in the mPFC, distinct from chronological age-related processes, emerges as a key determinant of individual variability in cognitive aging and offers a potential target for prevention and intervention.

A key insight from our work is that presynaptic mitochondria constitute a latent vulnerability that interacts with chronological age-related processes to cause cognitive decline, possibly through their ROS production, which is directly targeted by MitoQ. Although presynaptic mitochondrial abundance is not broadly altered by chronological age, it varies markedly among individuals even at young ages. Higher abundance predicts age-related cognitive decline; however, this relationship becomes evident only after aging. In aged mice, mPFC neurons show higher activity during attentional set shifting than in young mice, perhaps compensating for downregulation of glutamate receptors. This increased activity could enhance mitochondrial ROS via Ca^2+^ entry into mitochondria^36^. Moreover, because peroxisomes mitigate mitochondrial oxidative stress through contact-mediated ROS transfer^37^, age-associated downregulation of synaptic peroxisome functions, indicated by decreased peroxisomal proteins such as ABCD1-3 and ACOX1, could permit mitochondrial ROS to exert harmful effects. One of these effects could be impaired mitophagy, leading to the accumulation of ROS-producing mitochondria at synapses, because MitoQ treatment reduced presynaptic mitochondrial proteins. Thus, mitochondrial ROS and impaired mitophagy at the synapse may establish a vicious cycle, thereby contributing to progressive cognitive decline.

Although the precise mechanisms linking synaptic mitochondrial ROS to cognitive decline remain unresolved, synaptic proteomics with MitoQ implicate multiple pathways. For example, pro-apoptotic mitochondrial proteins, such as BAD, HTRA2, and BNIP3, may mediate ROS’s deleterious effects^38^, as MitoQ reduced their synaptosomal levels along with its pro-cognitive effect. Synaptic apoptosis signaling mediated by caspase-3 reportedly triggers microglial phagocytosis of synapses via complement cascades in cultured neurons^39^. In addition, since MitoQ upregulated several cytoskeletal regulators, including ADF/cofilin, mitochondrial ROS could also perturb synaptic actin dynamics, which are critical for synaptic function and plasticity^40^. Synaptic mitochondrial ROS may also suppress local protein translation, because mitochondrial oxidative stress activates the integrated stress response, which is known to reduce protein translation via eIF2α phosphorylation^41^. By contrast, in aged mice that maintain cognitive performance, this translational repression could be offset by higher levels of synaptic ribosomal proteins and mRNA-processing factors, including HNRNPC, a splicing protein that also regulates cytoplasmic mRNA metabolism and ribonucleoprotein assembly^42^.

In conclusion, synaptic mitochondria, particularly presynaptic, are critical determinants of individual variability in age-related cognitive decline. Their harmful influence likely reflects oxidative stress that arises when age-related factors, such as increased neuronal activity and peroxisome impairment, synergize with pre-existing vulnerabilities from youth, including defective clearance of dysfunctional mitochondria. This oxidative stress could then engage multiple downstream pathways, including release of mitochondrial pro-apoptotic proteins, suppression of local protein translation, and cytoskeletal reorganization, ultimately leading to cognitive decline. These synaptic mitochondria-centered mechanisms identified in this study offer a promising target for prevention and intervention for cognitive aging. Since dysfunctional mitochondria and impaired mitophagy are also implicated in age-related neurodegenerative diseases, including dementia, future studies should test whether synaptic mitochondrial oxidative stress also contributes to cognitive decline associated with these diseases.

## Acknowledgment

We thank Ando Akemi, Mikiko Suzuki, Misako Takizawa and Kaori Yamabe for secretarial help and Hiroko Iwamura for technical help. This study was supported in part by grants from AMED (JP25wm0625121 to H.N., JP24wm0425001, 25zf0127010, 25zf0127012, 25wm0625324 to T.F.), grants from JST Moonshot R&D (JPMJMS239F), Grant-in-Aid for Transformative Research Areas (23H04234 to T.F.) and Leading Initiative for Excellent Young Researchers (LEADER to H.N.) from the Ministry of Education, Culture, Sports, Science and Technology in Japan, Grants-in-Aid for Scientific Research (21H04812, 24K22086 to T.F., 20K07288, 23K06358 to H.N.) from the Japan Society for the Promotion of Science in Japan, the Cooperative Study Program of the National Institute for Physiological Sciences, and research grants from the Uehara Memorial Foundation (H.N.), Japan Foundation for Applied Enzymology (H.N.), the KANAE foundation for the promotion of medical science (H.N.), SENSHIN Medical Research Foundation (H.N., T.F.), Meiji Yasuda Life Foundation of Health and Welfare (H.N.), Daiichi Sankyo Foundation of Life Science (T.F.), SRF (T.F.), and the Kazato Foundation (H.N.).

## Author contributions

R.Y., H.N. and T.F. designed the study; R.Y., H.N., C.N., Y.Z., M.N., K.O., Y.K., N.O., T.F. performed and analyzed the results; H.N. and T.F. wrote the manuscript.

## Data availability

The datasets generated during and/or analyzed during the current study are available from the corresponding author on reasonable request.

## Notes

### Competing Interest Statement

The authors have declared no competing interest.

