## Supplementary figures for "Synaptic mitochondrial oxidative stress drives individual variability in age-related cognitive decline in mice"

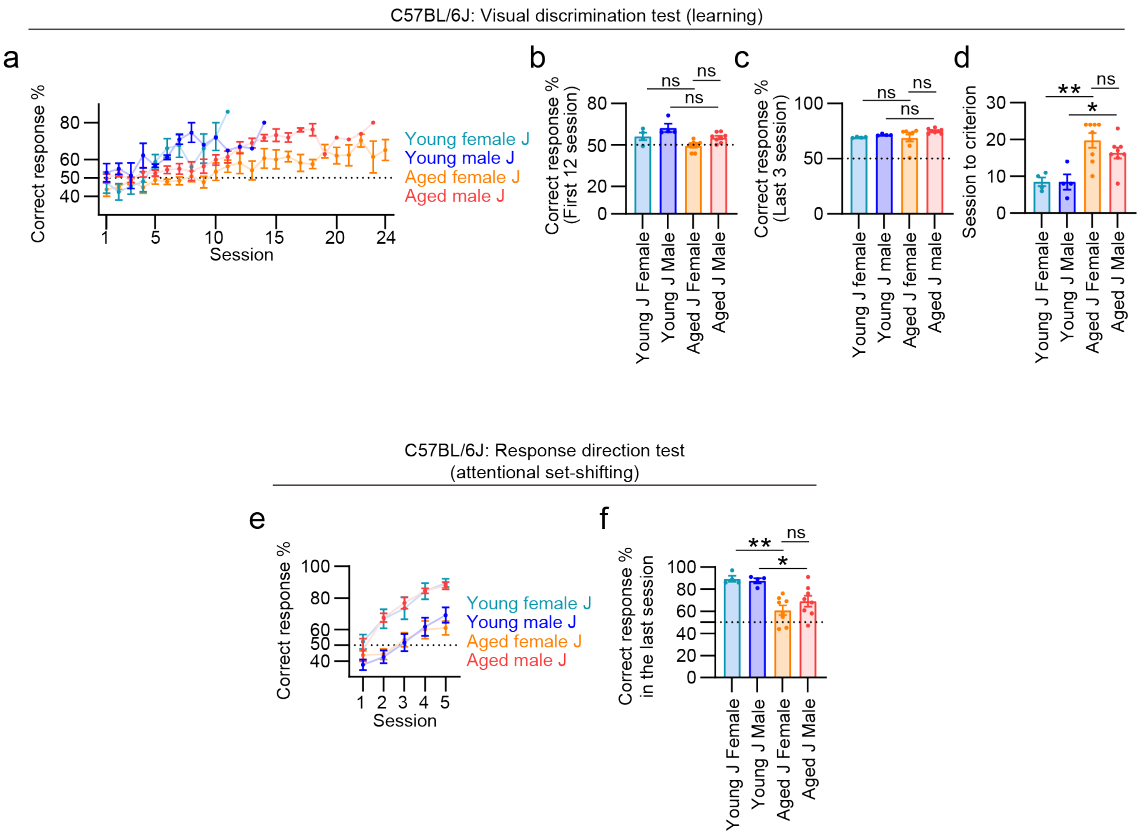


**Extended Data Fig. 1 Female and male C57BL/6J mice show similar age-related cognitive decline.** Behavioral performance of young and aged, male and female C57BL/6J mice in the visual discrimination and response direction tests. For the visual discrimination test, the correct response rate averaged within each session throughout the experiment (a), over the first 12 sessions (b), and over the last 3 sessions (c), as well as the number of sessions required to reach the performance criterion (d), are shown. For the response direction test, the correct response rate within each trial throughout the experiment (e) and within the last session (f) are shown. Data are presented as mean ± SEM. Statistical analyses were performed using one-way ANOVA with Holm-Sidak post hoc multiple comparisons. *$P$ < 0.05, **$P$ < 0.01; ns, not significant. Four young female, four young male, eight aged female, and eight aged male C57BL/6J mice were used.

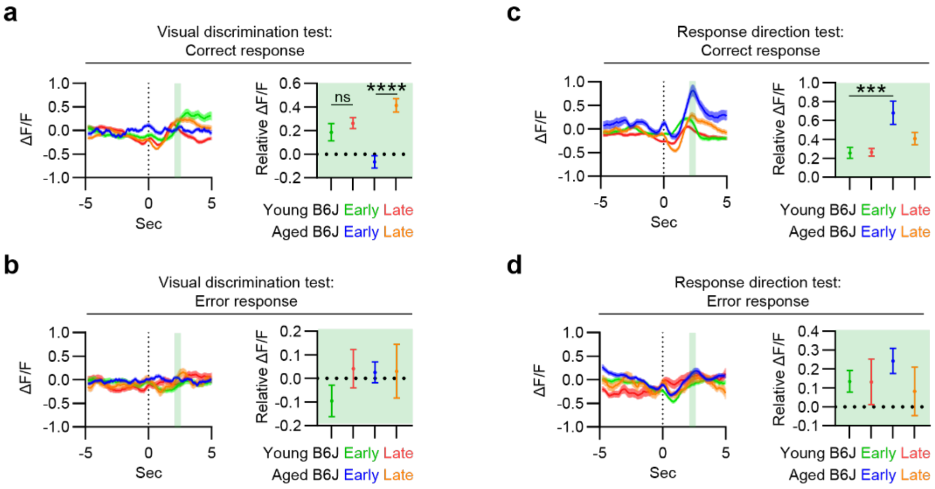


**Extended Data Fig. 2 mPFC neurons in aged mice show robust activation after correct behavioral responses in the visual discrimination and response direction tests.** The activities of mPFC neurons of young and aged male C57BL/6J mice, aligned to the timing of correct (a,c) or error (b,d) behavioral responses (time 0), in the visual discrimination test (a,b) and the response direction test (c,d) are shown in peri-event activity graphs. Relative ΔF/F values were calculated by averaging ΔF/F within predefined post-event time windows (2 to 2.5 s, indicated by the green background in the peri-event activity graphs) and subtracting the average ΔF/F around time 0 (-0.25 to 0.25 s). Data are presented as mean ± SEM. Statistical analyses were performed using one-way ANOVA with Holm-Sidak post hoc multiple comparisons. ***$P < 0.001$, ****$P < 0.0001$; ns, not significant. Four young and three aged male C57BL/6J mice were used.
